# Bacterial expression and purification of functional recombinant SARS-CoV-2 spike receptor binding domain

**DOI:** 10.1101/2021.02.03.429601

**Authors:** Janani Prahlad, Lucas R. Struble, William E. Lutz, Savanna A. Wallin, Surender Khurana, Andy Schnaubelt, Mara J. Broadhurst, Kenneth W. Bayles, Gloria E. O. Borgstahl

**Author notes:** To whom correspondence should be addressed: Gloria Borgstahl: University of Nebraska Medical Center, 986805 Nebraska Medical Center, Omaha, NE 68198-6805.

## Abstract

The COVID-19 pandemic caused by SARS-CoV-2 has applied significant pressure on overtaxed healthcare around the world, underscoring the urgent need for rapid diagnosis and treatment. We have developed a bacterial strategy for the expression and purification of the SARS-CoV-2 spike protein receptor binding domain using the CyDisCo system to create and maintain the correct disulfide bonds for protein integrity and functionality. We show that it is possible to quickly and inexpensively produce functional, active antigen in bacteria capable of recognizing and binding to the ACE2 (angiotensin-converting enzyme) receptor as well as antibodies in COVID-19 patient sera.

## Introduction

The Coronavirus disease-2019 (COVID-19) pandemic caused by the Severe Acute Respiratory Syndrome coronavirus 2 (SARS-CoV-2) has, due to its prolific interhuman transmission, become a dire global health concern (1). Since its discovery in Wuhan, China (2) in late 2019, there have been over 100 million cases worldwide, and over 2 million deaths as of January 2021. The novel betacoronavirus, SARS-CoV-2, belongs to the *Coronaviridae* family and is closely related to SARS-CoV-1 (79% genomic sequence identity) and MERS-CoV (Middle East respiratory syndrome coronavirus; 50% genomic sequence identity), two other pathogens responsible for the SARS epidemic of 2002, and the MERS epidemic in 2012, respectively (3,4). SARS-CoV-2 infection elicits a range of clinical presentations, from asymptomatic infection to severe viral pneumonia and death (3,4).

Like other coronaviruses, SARS-CoV-2 is comprised of four structural proteins: spike (S), envelope (E), membrane (M), and the nucleocapsid (N) (3,5,6). The S-protein is heavily glycosylated and can be found covering the surface of SARS-CoV-2 (7); glycosylation also allows the virus to evade the host immune system and gain entry into host cells via attachment to its receptor, ACE2 (5,7,8). At the amino acid level SARS-CoV-2 shares 90% identity with SARS-CoV-1(3). The S protein forms a homotrimeric class I fusion protein with each S monomer containing two subunits, S1 and S2 (4,5). When fused to host cell membranes, the S protein undergoes extensive conformational changes that cause dissociation of the S1 subunit from the complex and the formation of a stable post-fusion conformation of the S2 subunit (4,5). Much like SARS-CoV-1, receptor binding of the SARS-CoV-2 S protein relies on the receptor binding domain (RBD), which recognizes the aminopeptidase N segment of ACE2 (7). In addition to ACE2 recognition, the RBD is also responsible for eliciting neutralizing antibodies and has become a highly-investigated target for vaccine and drug development (3,5,6).

Spike RBD interaction with host ACE2 is the critical event preceding viral infection. Recent studies have shown that the RBD binds human ACE2 (hACE2) in low nanomolar affinity, further underscoring the importance of RBD in establishing viral attachment (4,5). The crystal structure of SARS-Cov-2 RBD has been described by Lan et al. (4) and highlights the key regions of the RBD responsible for ACE2 binding. The RBD consists of extensive β-sheets among which is nestled an extended insertion dubbed the receptor-binding motif (RBM); the RBM contains the residues that make contact with ACE2. The RBD also has ten cysteines (Fig. 1) which stabilize the overall structure through the formation of five disulfide bonds: three within the β-sheet core (C336-C361, C379-C432, and C391-C525), one connecting the distal loops of the RBM (C480-C488), and one that ties the N- and C-termini together (Cys538-Cys590) (4,5). It is important to note that not all of these disulfide bonds are consecutively formed and some require editing by reduction and reoxidation to form the correct bonds, which poses a complicated folding problem in the cytoplasm of bacteria (5).

**Fig 1.**
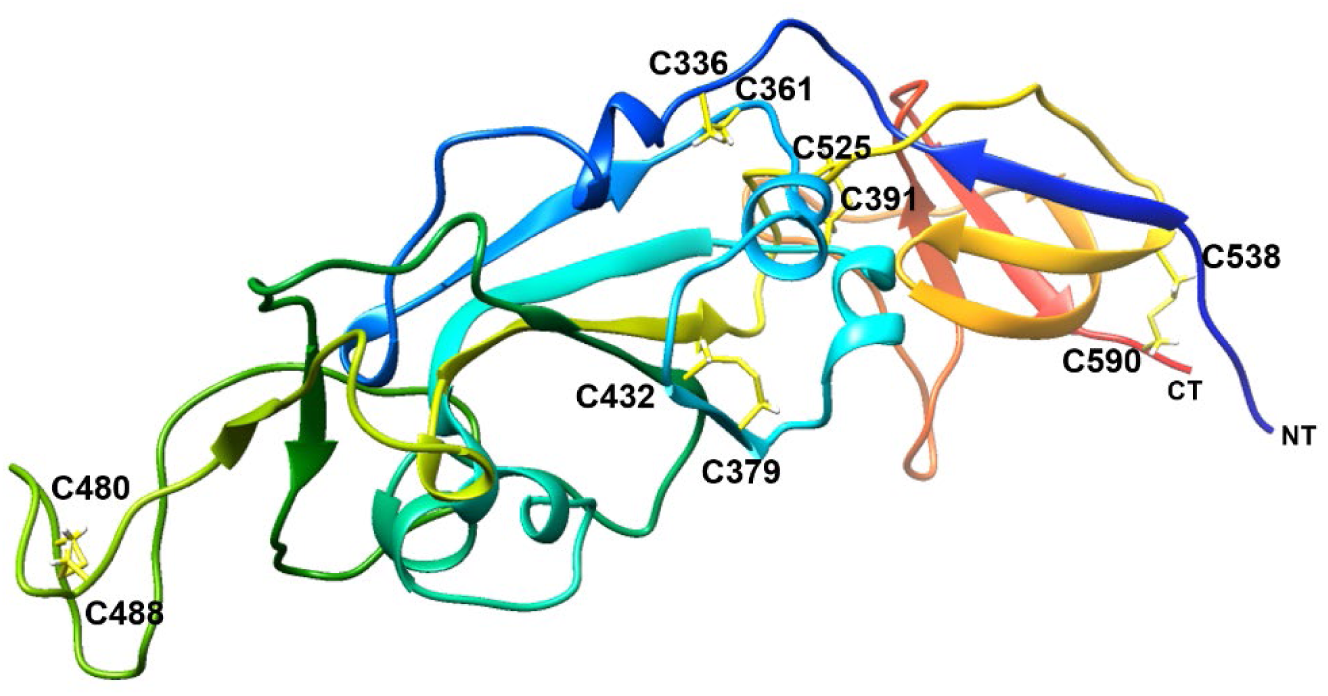
Ribbon diagram of SARS-CoV-2 spike RBD showing intramolecular disulfide bonds in yellow. As the structures solved for the RBD had missing gaps in the loops, we fed our sequence into I-Tasser, which generated the complete model (14-16). The ribbon was drawn using UCSF Chimera (17), is colored from blue (N-terminus) to red (C-terminus), and cysteines forming disulfide bonds are labeled..

Although advantageous in terms of ease of protein production, rapid growth, and cost-efficacy, the production of recombinant proteins in *E. coli* has its challenges (9,10). It is vital to maintain the correct disulfide bonds for structural as well as functional stability, which is problematic when expressing proteins of interest in the cytoplasm of wild-type *E. coli* (11). Disulfide bonds are typically formed in proteins which are secreted, or targeted to the outer membrane. The cellular organelles evolved to carry out this post-translational modification are the mitochondria and endoplasmic reticulum of eukaryotes, and the periplasmic space in prokaryotes (12). Disulfide bonds formed in the cytoplasm would be quickly reduced by multiple reductases and/or by reductants (such as glutaredoxin and thioredoxin). A possible solution would be to modify the protein construct to allow its secretion into the periplasm, which contains disulfide bond-catalyzing enzymes (9,11). However, this route greatly lessens the protein yield due to the limited cellular space occupied by the periplasm (8-16%), and requires modification of the construct to include a signal sequence for periplasmic targeting (9). Commercially available strains have been engineered to lack thioredoxin reductase and glutathione reductases, and to express disulfide bond isomerases, such as SHuffle (NEB), and Origami (Novagen), though these strains can suffer from low protein solubility and yield due to the lack of *de novo* or inappropriate disulfide bond formation (11).

To circumvent the drawbacks listed above, the Ruddock lab developed the CyDisCo (cytoplasmic disulfide bond formation in *E. coli*) system and showed that it was possible to produce active disulfide-bonded proteins in the cytoplasm of *E. coli* without deleting reduction systems, simply by expressing the sulfhydryl oxidase Evr1p (12,13). Evr1p uses molecular oxygen to form disulfides in a FAD-dependent reaction, allowing the production of disulfide-bonded proteins in the cytoplasm of wild-type *E. coli*. Another enzyme, protein disulfide isomerase (PDI), is added to edit disulfide bonds during protein folding. PDI can distinguish between properly folded and misfolded proteins, and correct disulfide bonds through cycles of cleavage and formation. In this way, correctly folded proteins with disulfide bonds can be recombinantly produced in the cytoplasm of *E. coli*.

We used the CyDisCo to make recombinant SARS-CoV-2 spike RBD properly folded with correct disulfide bonds. By cotransforming the spike RBD expression plasmid with the CyDisCo system, we were able to successfully produce recombinant spike RBD with the correctly-formed disulfide bonds. Here, we describe a simple cost-effective method of generating antigens using a bacterial expression system. By co-expressing our protein of interest along with the CyDisCo system, we have been able to produce correctly-folded antigen that retains activity and which can be used in diagnostic assays.

### Experimental Procedures

#### Protein expression and purification

The genetic sequence of spike RBD was fused to 10X-His-tagged maltose-binding protein (MBP), cloned into pET28a, and ordered from GenScript (Fig. 2) with codon-optimization for expression in *E. coli*. To ensure that all five disulfide bonds in the RBD (Fig. 1) (4) remain oxidized when expressed in the highly reducing cytoplasm of *E. coli*, we co-expressed the RBD-MBP fusion protein with a CyDisCo plasmid (9). The plasmids were co-expressed in one-shot BL21(DE3) cells (Thermofisher) in Luria-Bertani media (Fisher Bioreagents) under kanamycin (30 µg/ml) and chloramphenicol (34 µg/ml) selection. Cells were grown at 30° C, 275 rpm shaking until an OD_600_ of 0.3-0.4 was reached, and then induced with a final concentration of 0.5 mM isopropyl β-D-1-thiogalactopyranoside (IPTG) (Gold Biotechnology). At this point, the cells were cooled and transferred to 18° C, 150 rpm shaking overnight (approximately 16 hours) (Fig. 3). Cells were pelleted and stored at −20° C.

**Fig 2.**
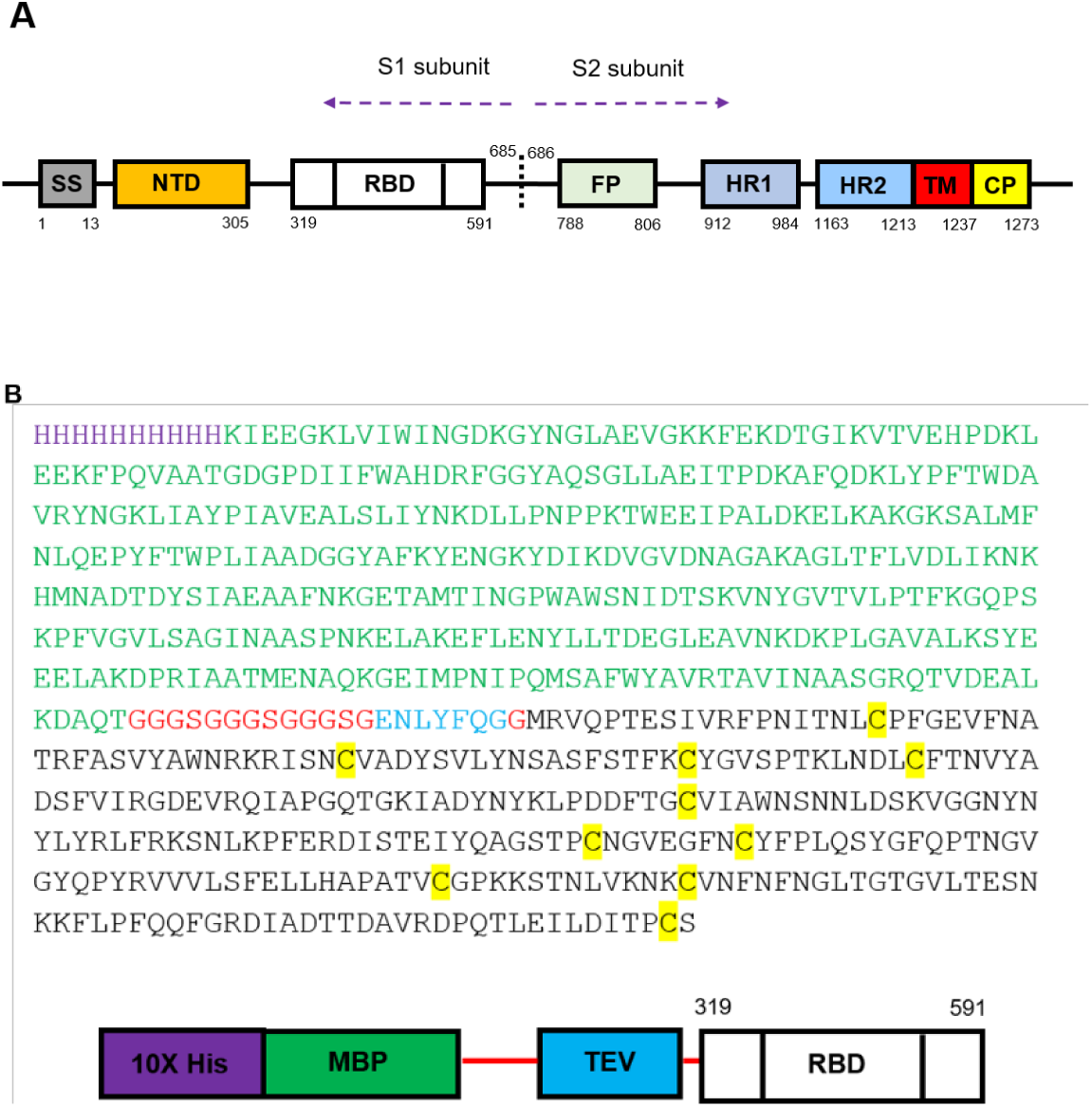
A) Schematic of full-length Spike protein SS: signal sequence; NTD: N-terminal domain; RBD: Ribosome-binding domain; FP: fusion peptide; HR1, HR2: heptad repeats 1 and 2; TM: transmembrane domain; CP: cytoplasmic peptide. B) Amino acid sequence of the RBD-MBP fusion protein used in this study.

**Fig 3.**
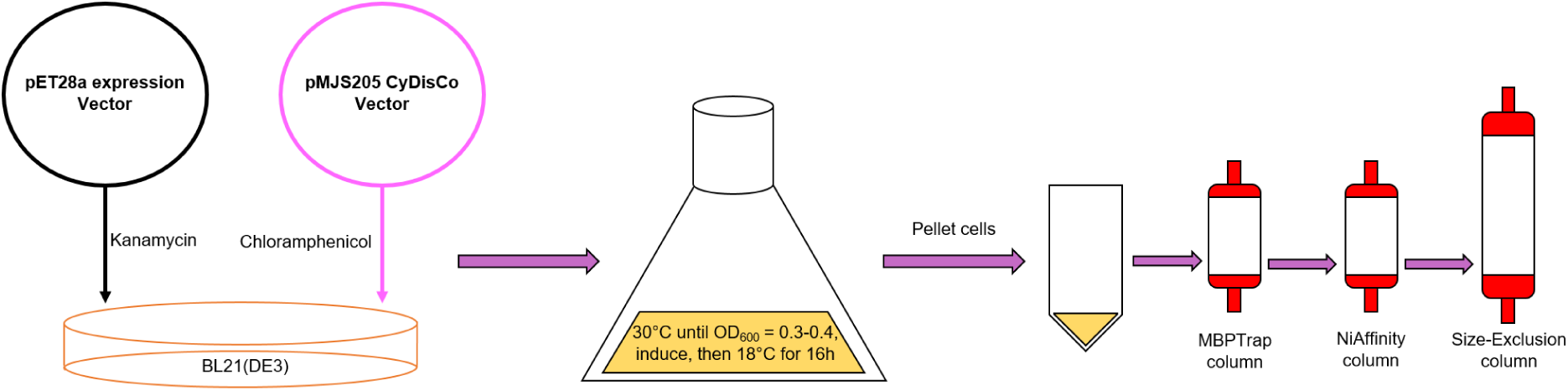
Experimental design for RBD-MBP expression and purification. RBD-MBP was cloned into pET28a and co-expressed with CyDisCo vector pMJS205 under kanamycin and chloramphenicol selection. A single colony of BL21(DE3) was inoculated into a few ml of media under antibiotic selection and scaled up into large two-liter flasks. Cells were grown at 30° C until the OD_600_ reached 0.3-0.4, at which point the cells were induced with 0.5 mM IPTG. The cells expressed protein overnight at 18°C and pelleted the following day. The cell pellet was lysed as described in the main text and passed through three chromatography columns to ensure optimal purity.

A cell pellet was thawed and resuspended in a buffer containing 20 mM Tris-HCl pH 7.4, 0.2 M NaCl, and 50 µl of protease inhibitor cocktail (Sigma, Cat.# P8849) per gram of cell pellet. The resuspended pellet was then lysed by three passes through an Emulsiflex C3. The resulting lysate was clarified by centrifugation at 40,000xg for 30 minutes at 4° C. Clarified lysate was then filtered through a 0.45 µm filter (Millipore) before being passed through a 5 ml MBPTrap column (GE) using an AKTA FPLC (Amersham Biosciences). Nonspecifically bound proteins were washed from the column with 10 CV of resuspension buffer, and RBD-MBP was eluted over 10 CV in a gradient method, using a buffer containing 20 mM Tris-HCl pH 7.4, 0.2 M NaCl, and 10 mM maltose. Fractions yielding protein (as observed by A_280_ peaks on the AKTA FPLC (Amersham Biosciences) chromatogram) were gel-verified using a 10% SDS-PAGE gel (Genscript) that was stained with Coomassie Brilliant Blue G-250 for size (Fig. 4), then loaded onto a 1 ml NiAff column (GE) with a running buffer containing 2X PBS with 20 mM imidazole, pH 8, and an elution buffer containing 2X PBS with 1 M imidazole, pH 8 over 16 CV. The resulting peaks were again gel-verified (Fig. 4), and lanes with bands corresponding to 74 kDa (expected size of the RBD-MBP fusion protein) were concentrated and loaded onto a HiLoad 16/60 Superdex75 (GE) size-exclusion column. Fractions corresponding to the fusion protein (Fig. 4), were pooled and concentrated (using Amicon regenerated cellulose concentrators (Cat # UFC803096). RBD-MBP identity after each column was confirmed by Peggy Sue (Protein Simple) western analysis using Sino Biological Anti-Coronavirus spike antibody (Cat # 40591-T62). The concentrations were determined using a Fisher NanoDrop1000 using a molecular weight of 74 kDa for the fusion protein, and calculated ε_280_=101.19 M^−1^cm^−1^. The yield from 2 liters of culture was approximately 0.5 mg purified RBD-MBP. Purified RBD-MBP was stored at −20°C in 2X PBS supplemented with a final concentration of 30% glycerol.

**Fig 4.**
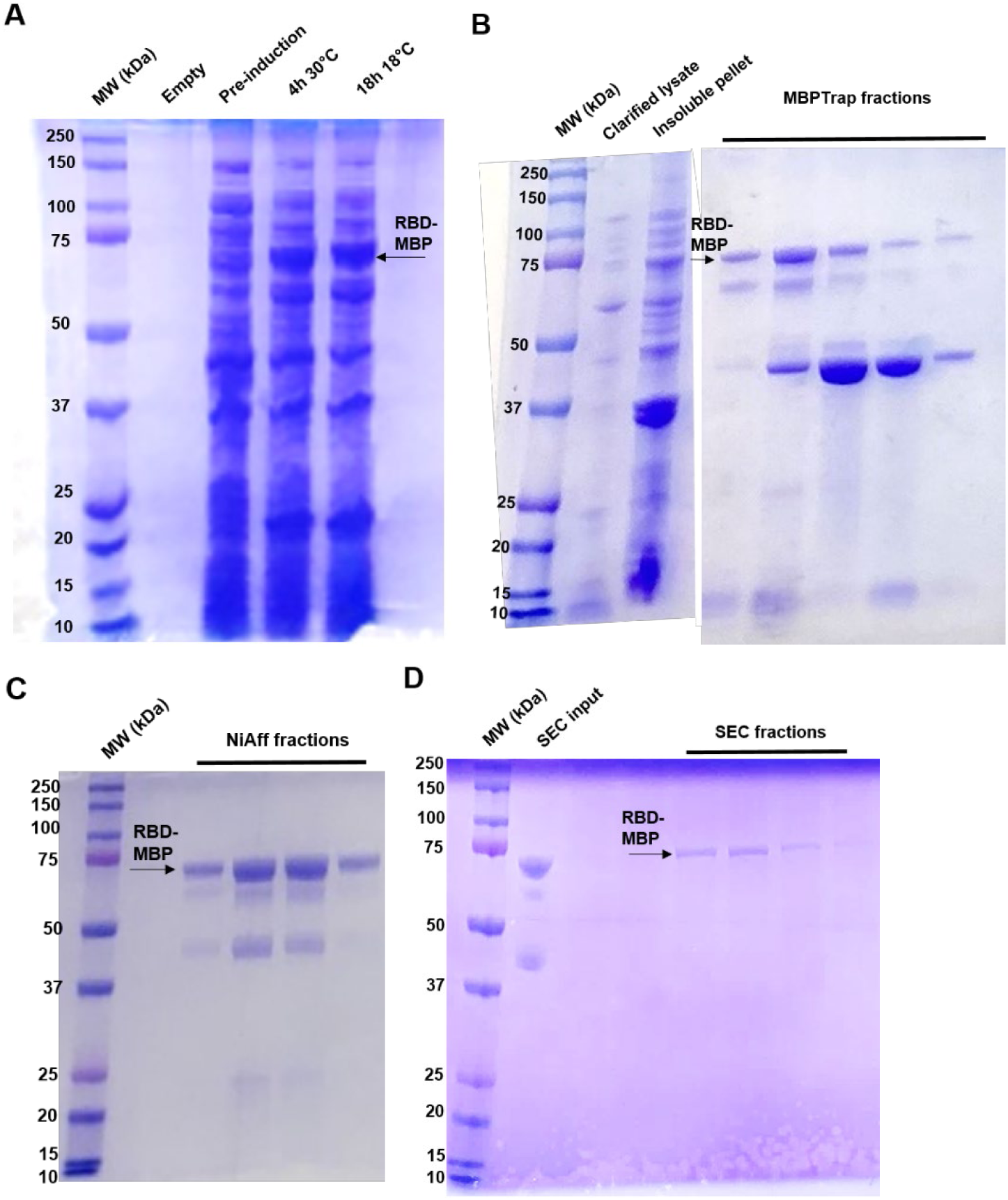
Protein purification SDS-PAGE gels. RBD-MBP can be observed running at 74 kDa, as indicated by the black arrows near the 75 kDa marker. Black arrows at approximately 42 kDa could indicate endogenously produced or cleaved MBP. A) Initial induction experiment comparing same-day (4 h) induction at 30° C and overnight (16 h) induction at 18° C. B) MBPTrap AKTA fractions. C) Nickel-affinity AKTA fractions. D) Size-exclusion AKTA fractions.

#### Surface Plasmon Resonance (SPR) based hACE2 binding assay

The recombinant hACE2-AviTag protein from 293T cells (Acro Biosystems, DE, USA) was captured on a sensor chip in the test flow channels. Samples of 300 μl of freshly prepared serial dilutions of the purified recombinant protein were injected at a flow rate of 50 μl/min (contact duration 180 seconds) for the association. Responses from the protein surface were corrected for the response from a mock surface and for responses from a buffer-only injection. Total hACE2 binding and data analysis were calculated with Bio-Rad ProteOn Manager software (version 3.1).

#### Luminex serological assay

This assay was performed using the purified SARS-CoV-2 RBD-MBP fusion (5 µg per 106 beads) coupled to the surface of group A, region 43, MagPlex® Microspheres (Luminex Corp, IL, USA). Microsphere coupling was performed using the Luminex xMAP Antibody Coupling Kit (Luminex Corp, IL, USA) according to the manufacturer’s instructions. The protein-coupled microspheres were re-suspended in PBS-TBN buffer (1X PBS containing 0.1% Tween 20, 0.5% BSA and 0.1% sodium azide) at a final stock concentration of 2×10^6^ microspheres per ml, with 2.5×10^3^ microspheres used per reaction. De-identified serum samples were obtained from residual patient sera that tested positive or negative by a clinical SARS-CoV-2 serology assay (Diasorin LIASON SARS-CoV-2 S1/S1 IgG; n=6 per group) at UNMC. Fifty microliters of each serum sample (diluted 1:50 with 1X PBS-TBN buffer) was mixed with 50 μl of the RBD-MBP coupled microspheres in a 96-well plate. The assay plate was incubated for 30 minutes at 37° C with shaking at 700 rpm and then washed 5 times with 1X PBS-TBN buffer. Then, the plate was incubated with biotin-conjugated goat anti-human IgG (Abcam, MA, USA), labeled with streptavidin R-phycoerythrin reporter (Luminex xTAG® SA-PE G75) for 1 hour at 25° C with shaking at 700 rpm. Then, the plate was washed 5 times with 1X PBS-TBN buffer. Finally, the plate was re-suspended in 100 ml of 1X PBS-TBN buffer and incubated for 10 minutes at 25°C with shaking at 700 rpm. The microplate was assayed on a Luminex MAGPIXTM System, and results were reported as median fluorescent intensity (MFI).

## Results and Discussion

### Purification of Spike RBD-MBP fusion protein

The gene sequence for the Spike RBD-MBP fusion protein was cloned into pET28a and codon-optimized by Genscript. To facilitate cleavage of MBP from RBD, a TEV recognition sequence was included after the MBP protein sequence. RBD-MBP purified readily from BL21(DE3) *E. coli* pellets. Initial IPTG induction tests showed appreciable accumulation of the fusion protein after 16-18 hours at 18° C (Fig. 4). Though induction appeared similar to 4 hour induction at 30° C, it was decided that lower-temperature induction for a longer time would encourage thorough editing and formation of disulfide bonds and correct folding of the recombinant protein. The pellets were thawed and lysed, in the presence of protease inhibitor and all steps were performed with the lysate on ice or at 4° C. After the lysate was clarified by centrifugation, it was passed through a MBPTrap column to clear non-specific proteins lacking the MBP tag. Fractions containing RBD-MBP were confirmed through SDS-PAGE (Fig. 4) where RBD-MBP can be observed running at ∼74 kDa (indicated by red arrows). Interestingly, we could also see prominent bands at ∼42 kDa (indicated by the green arrow), corresponding to the size of MBP, indicating that we also pulled down endogenously-produced *E. coli* MBP. The resulting fractions were pooled before passing through a NiAff column, which bound the 10X His tag on the fusion protein. This additional affinity purification step helped reduce the amount of non-specific proteins that co-purified with RBD-MBP, as well as reduce the presence of any cleaved or endogenous MBP from the pool of fusion protein. Finally, RBD-MBP was further purified using a size-exclusion step over a Superdex75 column. Using the steps outlined above, we were able to obtain approximately 0.5 mg of pure RBD-MBP from 2 liters of culture.

### Binding of Spike RBD-MBP to hACE2 receptor

To test the activity of purified RBD-MBP, we assessed its binding to hACE2 using SPR against control RBD secreted from human cells (Fig. 5). We showed that the purified RBD-MBP bound hACE2, indicating correct functional folding of the purified recombinant protein.

**Fig 5.**
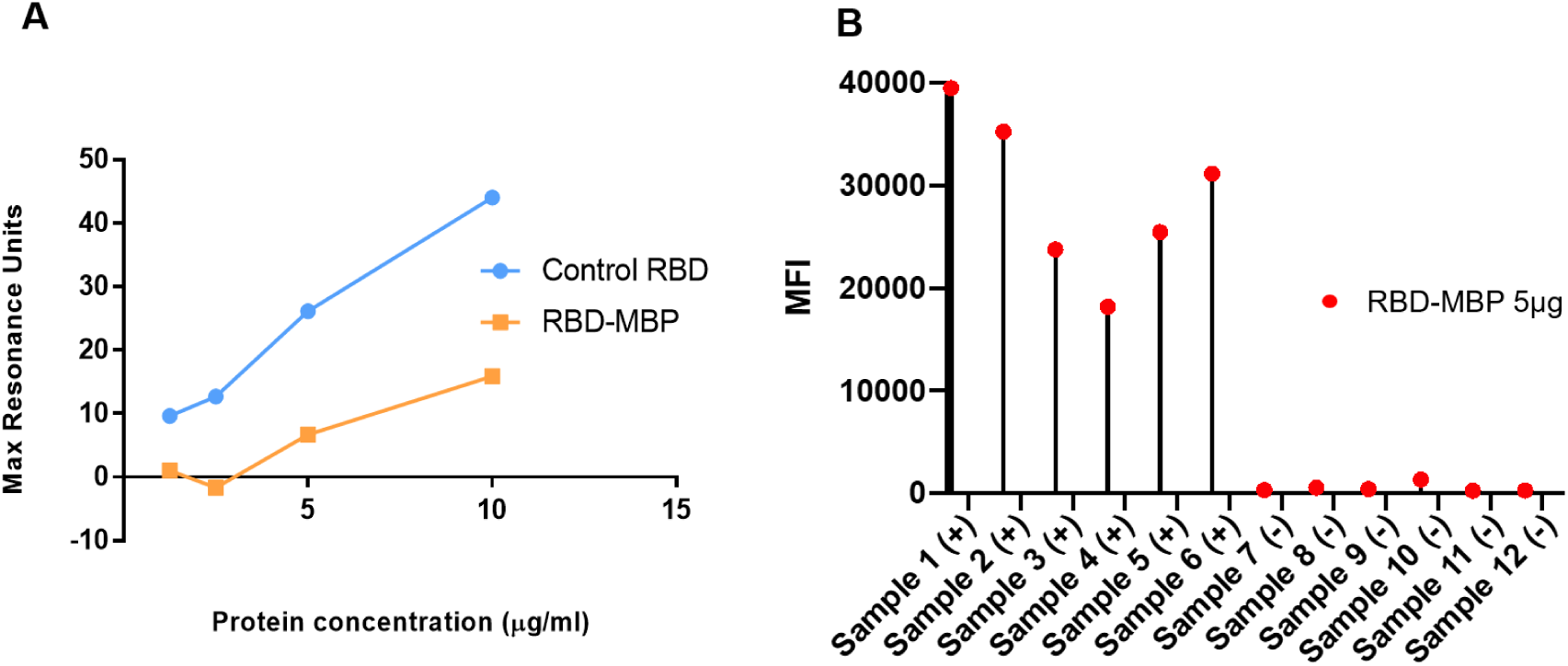
Activity assessment of purified recombinant RBD-MBP. A) SPR binding assay of RBD-MBP to hACE2. B) Microsphere immunoassay of RBD-MBP binding to IgG antibodies in patient sera [(+) = SARS-CoV-2 IgG positive, (-) = SARS-CoV-2 IgG negative]. MFI: mean fluorescence intensity.

### SARS-CoV-2 antibodies in patient serum samples bind to Spike RBD-MBP

An immunoassay was performed to confirm that our RBD-MBP was recognized by human SARS-CoV-2-specific antibodies. RBD-MBP-bound microspheres were incubated with sera that had tested positive (n=6) or negative (n=6) in a clinical SARS-CoV-2 serology assay. RBD-specific IgG antibodies were detected in all sera from SARS-CoV-2 antibody-positive patients (Fig. 5).

### Conclusions

In this paper, we have described a rapid method to express and purify functionally active RBD antigen in E. coli. The co-expression of the sulfhydryl oxidase, Evr1p, and the disulfide bond isomerase, PDI in the CyDisCo system along with the plasmid of interest results in a well-folded and functionally active protein in a bacterial system, which is simple, rapid, and less expensive than a eukaryotic or mammalian system. It is noteworthy that the RBD expressed in bacteria here does not have any post-translational modifications.

## Acknowledgments

The authors thank Dr. Lloyd Ruddock at the University of Oulu, Oulu, Finland for kindly gifting the CyDisCo plasmid and providing essential advice on its use for this study. This research was supported by the Fred and Pamela Buffett NCI Cancer Center Support Grant (P30CA036727) and its associated COVID19 supplement.

